# Characterizing the mechanical properties of low-profile prosthetic feet

**DOI:** 10.1101/2023.04.14.536964

**Authors:** Joshua R. Tacca, Zane A. Colvin, Alena M. Grabowski

**Affiliations:** Paul M. Rady Department of Mechanical Engineering, University of Colorado, Boulder, CO, USA; Department of Integrative Physiology, University of Colorado, Boulder, CO, USA; Department of Veterans Affairs, Eastern Colorado Healthcare System, Denver, CO, USA

**Keywords:** prostheses, stiffness, hysteresis, amputation

## Abstract

Passive-elastic prosthetic feet are manufactured with different numerical stiffness categories that are prescribed based on the body mass and activity level of the user, but the mechanical properties, such as the stiffness values and hysteresis are not typically reported by the manufacturer. Since the mechanical properties of passive-elastic prosthetic feet can affect the walking biomechanics of people with transtibial or transfemoral amputation, characterizing these properties would provide objective values for comparison and aid the prescription of prosthetic feet. Therefore, we characterized the axial stiffness values, torsional stiffness values, and hysteresis of 33 different categories and sizes of a commercially available passive-elastic prosthetic foot model, the Össur low-profile (LP) Vari-flex with and without a shoe. We measured axial stiffness from compression and torsional stiffness from dorsiflexing and plantarflexing the prostheses. In general, a greater numerical prosthetic foot stiffness category resulted in increased heel, midfoot, and forefoot axial stiffness values, increased plantarflexion and dorsiflexion torsional stiffness values, and decreased heel, midfoot, and forefoot hysteresis. Moreover, a greater prosthetic foot size resulted in decreased heel, midfoot, and forefoot axial stiffness values, increased plantarflexion and dorsiflexion torsional stiffness values, and decreased heel and midfoot hysteresis. Finally, adding a shoe to the LP Vari-flex prosthetic foot resulted in decreased heel and midfoot axial stiffness values, decreased plantarflexion torsional stiffness values, and increased heel, midfoot, and forefoot hysteresis. In addition, we found that the force-displacement and torque-angle relationships were better described by curvilinear than linear equations. Ultimately, our results can be used to objectively compare LP Vari-flex prosthetic feet to other prosthetic feet in order to inform their prescription and design and use by people with a transtibial or transfemoral amputation.

## 1. Introduction

To walk, people with a transtibial or transfemoral amputation typically use passive-elastic prosthetic feet, which are comprised of carbon fiber or fiberglass, and allow elastic energy storage and return during the stance phase. The mechanical properties of passive-elastic prosthetic feet, such as stiffness and hysteresis, affect kinematics, kinetics, muscle activity, metabolic cost and user preference during walking (1–7). Yet, these mechanical properties are not typically reported by the manufacturer. Instead, prosthetic manufacturers use numerical stiffness categories (e.g., 1-9) where a higher numerical stiffness category corresponds to a stiffer prosthesis and numerical stiffness categories are prescribed based on the body mass and activity level of the user (8). However, these stiffness categories and the differences in stiffness values between categories are not consistent across manufacturers (9,10). To better inform prosthetic foot prescription, objective values for mechanical properties of prosthetic feet such as stiffness values and hysteresis should be provided. These values will inform dynamic function and can be implemented in future prosthetic design.

Previous studies have characterized the mechanical properties of passive-elastic prosthetic feet (9–18) and found that they are well described by linear (11,14) or curvilinear (9,13,15–17) force-displacement and torque-angle profiles, which are used to calculate axial (kN/m) and torsional (kN*m/rad) stiffness values. These studies provide axial stiffness values for forces applied to a prosthetic foot heel, midfoot, and forefoot. The axial stiffness values of different portions of prosthetic feet can affect the biomechanics of the user during walking (1). For example, a study that varied the stiffness values of the heel and forefoot of an experimental passive-elastic prosthetic foot, found that greater heel stiffness values resulted in a higher ground reaction force loading rate, greater knee flexion angle in early stance, and greater knee extension moment, and that greater forefoot stiffness values resulted in greater knee extension angle in mid-stance and greater knee flexor moment during walking at a range of speeds (0.7 to 1.5 m/s) for people with unilateral transtibial amputation (1). Therefore, determining the axial stiffness values for the heel, midfoot, and forefoot of a passive-elastic prosthetic foot would provide objective values for comparison between prosthetic feet and better predict the effects of different passive-elastic prosthetic foot stiffness values on walking biomechanics. Moreover, since a biological ankle can behave mechanically like a torsional spring and damper system during walking at 1.2 m/s (19), the torsional stiffness values of passive-elastic prosthetic feet provide additional information that can be compared to a biological ankle-foot system to derive function. Some previous studies have characterized torsional prosthetic foot stiffness properties (6,14,20) and found that prosthetic foot length affects torsional stiffness values since longer feet have a longer moment arm and for a given applied force and angle change, have greater torsional stiffness values than shorter feet (20). Furthermore, the energy returned by a prosthetic foot during the push-off phase of stance depends on both stiffness values (3) and hysteresis, or energy loss (15,21). Energy return of a passive-elastic prosthetic foot is related to the energy stored and can affect walking biomechanics, where lower energy return can result in decreased affected leg work during push-off, increased unaffected leg work during collision, and increased hip work (7,22). Hysteresis has been reported for some passive-elastic prosthetic feet (11,15–17) and likely depends more on material properties rather than stiffness categories of prosthetic feet (11). Ultimately, characterizing the axial and torsional stiffness values and hysteresis of passive-elastic prosthetic feet will better inform prosthetic prescription and function by allowing objective comparisons between different prosthetic foot stiffness categories and sizes.

Most people with a transtibial or transfemoral amputation wear shoes over their prosthetic foot during walking and this likely affects overall stiffness values and hysteresis compared to prosthetic feet alone (23). Major et al. found that wearing shoes over prosthetic feet resulted in lower axial stiffness values at the heel and midfoot but not at the forefoot compared to prosthetic feet alone (23). Moreover, Major et al. found that wearing shoes over prosthetic feet resulted in increased hysteresis compared to prosthetic feet alone (23). Since adding shoes to prosthetic feet changes the overall mechanical properties compared to prosthetic feet alone and because shoes are commonly used when people with a transtibial or transfemoral amputation walk, shoes should be considered when characterizing the mechanical properties of prosthetic feet.

There are many prosthetic foot models that are commercially available to people with a transtibial or transfemoral amputation. One such model is the Össur low-profile (LP) Vari-flex (Össur, Reykjavik, Iceland), which is a passive-elastic prosthetic foot made of carbon-fiber with a short build height (0.068 m) (8) so that it can be used by people with long residual limbs or fit beneath a stance-phase powered ankle-foot prosthesis such as the BiOM (now Ottobock Empower, Duderstadt, Germany) (24). Characterizing the mechanical properties of LP Vari-flex feet will provide objective measures that can be used to compare these prostheses to other available prosthetic feet, inform dynamic function, and influence future prosthetic design. Therefore, we measured the axial and torsional stiffness values and hysteresis of different stiffness categories and sizes of LP Vari-flex feet. We measured axial stiffness from compression and torsional stiffness from dorsiflexing and plantarflexing the prostheses. Moreover, since shoes can affect the mechanical properties of prosthetic feet, we measured the axial and torsional stiffness values and hysteresis of LP Vari-flex feet with and without a representative walking shoe.

First, since the Össur Vari-flex (higher profile version of the LP Vari-flex, Össur, Reykjavik, Iceland) exhibits a curvilinear force-displacement profile (9), we hypothesized that the axial and torsional stiffness of all the stiffness categories of the LP Vari-flex would be better characterized by a curvilinear force-displacement profile than a linear force-displacement profile independent of a shoe. Second, we hypothesized that because a greater numerical stiffness category is prescribed to people with greater body mass and higher activity levels, a greater stiffness category would result in higher axial stiffness values when force is applied at the heel, midfoot, and forefoot, and higher torsional stiffness values when plantarflexion and dorsiflexion torque are applied to the prosthetic foot but would have no effect on hysteresis with or without a shoe. Third, we hypothesized that greater passive-elastic prosthetic foot length (size) would have no effect on axial stiffness values when force is applied at the heel, midfoot, and forefoot, and hysteresis within a given category but would result in greater torsional stiffness values when plantarflexion and dorsiflexion torque are applied to the prosthetic foot with or without a shoe. Fourth, we hypothesized that adding a shoe to the prosthetic foot would result in lower axial stiffness values when force is applied at the heel and midfoot and increase hysteresis but not affect axial stiffness values when force is applied at the forefoot or torsional stiffness values when plantarflexion and dorsiflexion torque are applied to the prosthetic foot and shoe compared to without a shoe.

## 2. Methods

### Prosthetic feet

LP Vari-flex prosthetic feet are manufactured in a range of different stiffness categories (1–9) and foot sizes (22-30 cm) that are prescribed to people with a transtibial or transfemoral amputation who have a range of body mass (45-166 kg) and low to high impact (activity) levels (8). We determined the axial stiffness (kN/m) values, torsional stiffness (N*m/rad) values, and hysteresis (%) of 33 different LP Vari-flex prosthetic feet with different stiffness categories (Categories 1-8) and sizes (24-29 cm; Table 1) in compression using a materials testing machine (MTM; Instron Series 5859, Norwood, MA). We determined axial stiffness values and hysteresis with a force applied at the heel, midfoot, and forefoot of each prosthetic foot including the rubber cosmesis with and without a standard walking shoe (New Balance MA411, Boston, MA). Then, we determined torsional stiffness values when plantarflexion and dorsiflexion torque were applied to each prosthetic foot including the rubber cosmesis with and without a standard walking shoe (New Balance MA411, Boston, MA). For two prosthetic sizes (27 and 29), we did not have a New Balance MA411 shoe, so we instead used New Balance MW928 shoes.

**Table 1.** Low-profile Vari-flex prosthetic foot (8) stiffness category, size, shoe, manufacturer recommended body mass range for a moderate impact level, 1.3 times the maximum recommended body weight (BW; maximum force threshold for the heel and midfoot tests), and 1.0 times the maximum recommended BW (maximum force threshold for the forefoot test).

| Foot | Stiffness Category | Size (cm) | Shoe (US Size) | Recommended Body Mass (kg) [Moderate Impact Level] | 1.3 BW (N) | 1.0 BW (N) |
| --- | --- | --- | --- | --- | --- | --- |
| 1 | 1 | 24 | MA411 (7) | 45–52 | 662 | 510 |
| 2 | 1 | 25 | MA411 (8) | 45–52 | 662 | 510 |
| 3 | 1 | 26 | MA411 (9) | 45–52 | 662 | 510 |
| 4 | 2 | 24 | MA411 (7) | 53–59 | 752 | 578 |
| 5 | 2 | 25 | MA411 (8) | 53–59 | 752 | 578 |
| 6 | 2 | 26 | MA411 (9) | 53–59 | 752 | 578 |
| 7 | 3 | 24 | MA411 (7) | 60–68 | 866 | 666 |
| 8 | 3 | 25 | MA411 (8) | 60–68 | 866 | 666 |
| 9 | 3 | 26 | MA411 (9) | 60–68 | 866 | 666 |
| 10 | 3 | 28 | MA411 (11) | 60–68 | 866 | 666 |
| 11 | 3 | 29 | MW928 (12.5) | 60–68 | 866 | 666 |
| 12 | 4 | 24 | MA411 (7) | 69–77 | 981 | 755 |
| 13 | 4 | 25 | MA411 (8) | 69–77 | 981 | 755 |
| 14 | 4 | 26 | MA411 (9) | 69–77 | 981 | 755 |
| 15 | 4 | 27 | MW928 (10) | 69–77 | 981 | 755 |
| 16 | 4 | 28 | MA411 (11) | 69–77 | 981 | 755 |
| 17 | 4 | 29 | MW928 (12.5) | 69–77 | 981 | 755 |
| 18 | 5 | 24 | MA411 (7) | 78–88 | 1121 | 862 |
| 19 | 5 | 25 | MA411 (8) | 78–88 | 1121 | 862 |
| 20 | 5 | 26 | MA411 (9) | 78–88 | 1121 | 862 |
| 21 | 5 | 27 | MW928 (10) | 78–88 | 1121 | 862 |
| 22 | 5 | 28 | MA411 (11) | 78–88 | 1121 | 862 |
| 23 | 5 | 29 | MW928 (12.5) | 78–88 | 1121 | 862 |
| 24 | 6 | 25 | MA411 (8) | 89–100 | 1274 | 980 |
| 25 | 6 | 26 | MA411 (9) | 89–100 | 1274 | 980 |
| 26 | 6 | 27 | MW928 (10) | 89–100 | 1274 | 980 |
| 27 | 6 | 28 | MA411 (11) | 89–100 | 1274 | 980 |
| 28 | 6 | 29 | MW928 (12.5) | 89–100 | 1274 | 980 |
| 29 | 7 | 26 | MA411 (9) | 101–116 | 1478 | 1137 |
| 30 | 7 | 27 | MW928 (10) | 101–116 | 1478 | 1137 |
| 31 | 7 | 28 | MA411 (11) | 101–116 | 1478 | 1137 |
| 32 | 7 | 29 | MW928 (12.5) | 101–116 | 1478 | 1137 |
| 33 | 8 | 27 | MW928 (10) | 117–130 | 1656 | 1274 |

### Axial Stiffness

We constructed a custom aluminum base and low-friction roller system for the MTM to measure the heel, midfoot, and forefoot axial stiffness values of the prosthetic feet. We used a low friction roller between the base and each prosthetic foot to minimize torque on the uniaxial load cell of the MTM (Fig. 1). A rigid pylon was aligned vertically and attached to the MTM. Each prosthetic foot was attached to the rigid pylon and the bottom of the prosthetic foot was aligned perpendicular to the pylon. We set the base at −15°, 0°, and 20° relative to horizontal, which corresponds to the angles required for heel, midfoot, and forefoot axial stiffness testing, respectively (25) (Fig. 1). For each test, we preloaded the prosthetic foot with 4-6 N and used the MTM to apply a force along the pylon at 100 N/s (25) for four consecutive loading and unloading cycles. Each prosthetic foot stiffness category is recommended for a user within a range of body mass values by the manufacturer (Table 1). For each prosthetic stiffness category, we used the highest body mass within the recommended range to estimate the ground reaction force applied on the heel, midfoot, and forefoot of the prosthetic foot during walking. We set the maximum force of each test to a value based off the first and second peak ground reaction forces on the affected leg of a person with a transtibial amputation walking on level ground at 1.75 m/s to estimate the ground reaction force that could be applied to a particular prosthesis during walking (26). When a person with a transtibial amputation walks on level ground at 1.75 m/s using a passive-elastic prosthesis, they apply a first peak vertical ground reaction force that is 1.3 times their bodyweight (BW) for their affected leg (26). Thus, we applied a maximum force of 1.3 times BW of the heaviest person within the recommended range for each prosthetic stiffness category at a moderate impact level for the heel and midfoot tests (base at −15° and 0°). When a person with a transtibial amputation walks on level ground at 1.75 m/s using a passive-elastic prosthesis, they apply a second peak vertical ground reaction force that is 1.0 times their bodyweight (BW) for their affected leg (26). Thus, we applied a maximum force of 1.0 times BW of the heaviest person within the recommended range for each prosthetic stiffness category at a moderate impact level for the forefoot test (base at 20°).

**Fig. 1.**
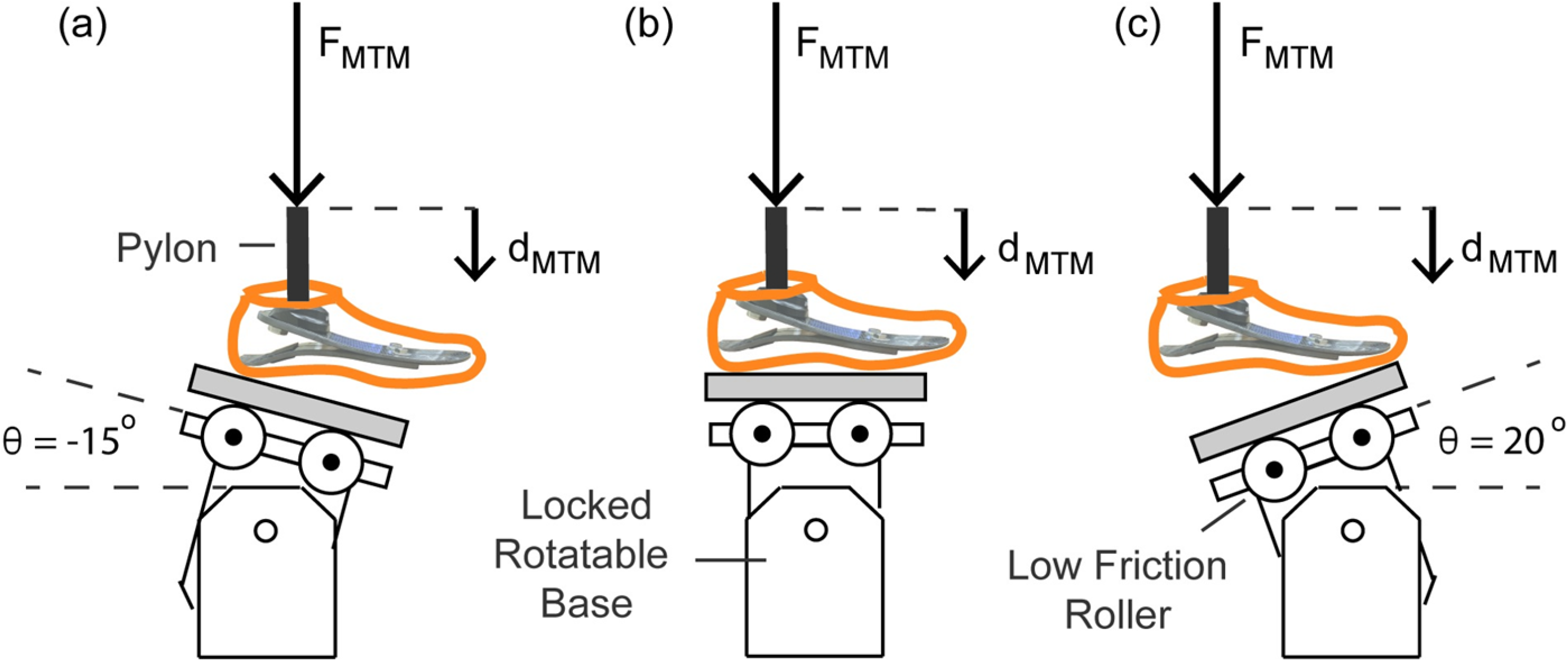
**Illustration of axial stiffness testing for the (a) heel, (b) midfoot, and (c) forefoot of each prosthetic foot.** The materials testing machine (MTM) applied force (F_MTM_) vertically at 100 N/s along the pylon to compress the prosthetic foot. F_MTM_ and vertical displacement (d_MTM_) were measured by the load cell and MTM. A low friction roller was placed beneath the prosthetic foot to minimize the torque applied on the load cell of the MTM. For the axial tests on the heel, midfoot, and forefoot, the rotatable base was locked at −15°, 0°, and 20° relative to horizontal, respectively.

We determined the axial stiffness values of each prosthetic foot as the quotient of the normal force and displacement applied by the base onto the bottom of the prosthesis (Fig. 1, 2). The normal force (F_norm_) equals the quotient of the F_MTM_ and the cosine of the angle of the base relative to the prosthetic foot (θ) (Fig. 1, 2):

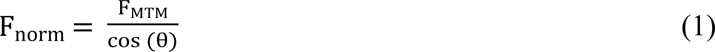

**Fig. 2.**
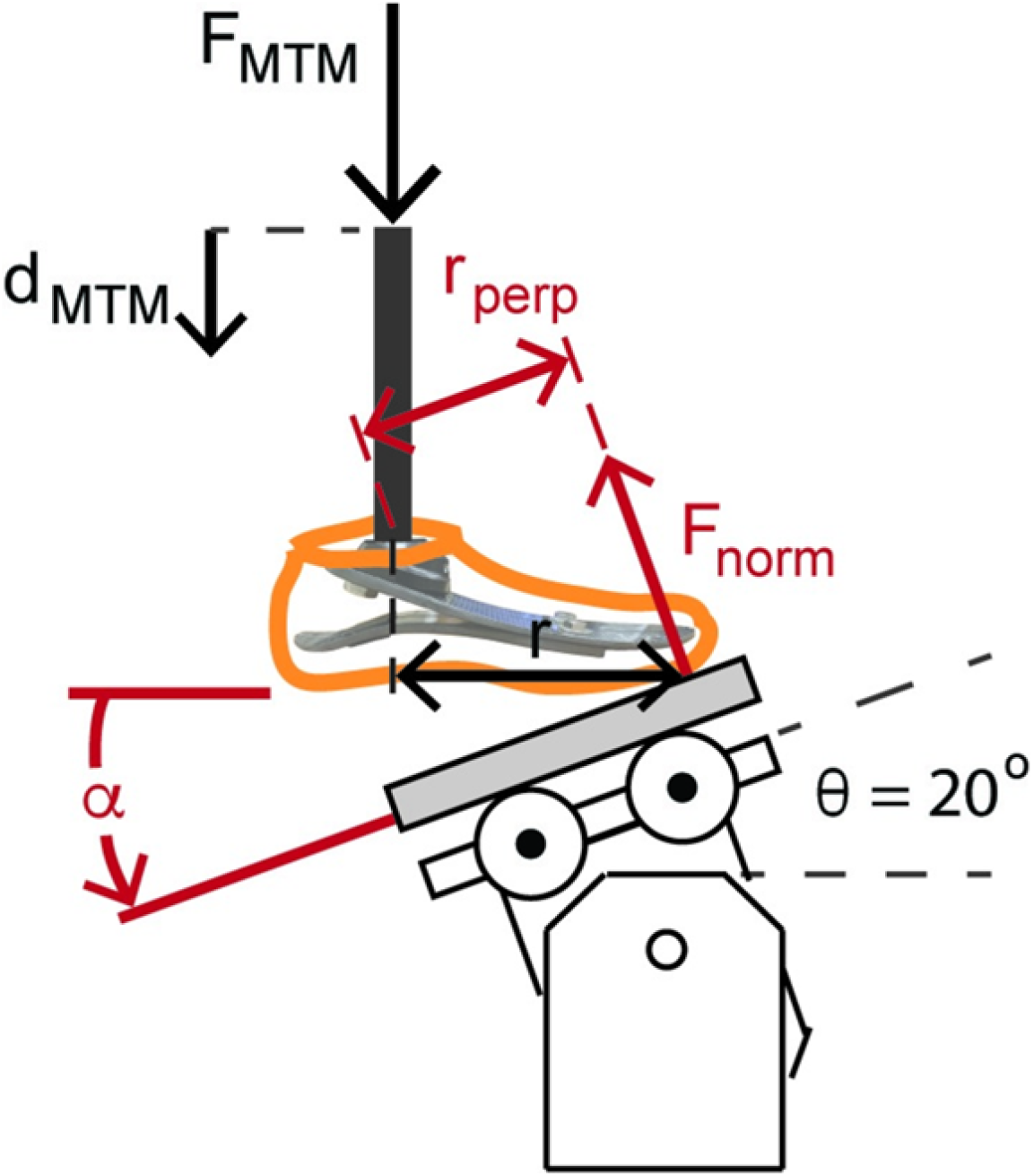
**Illustration of the forefoot prosthetic axial stiffness testing and dorsiflexion torsional stiffness testing.** The MTM applied force (F_MTM_) and displacement (d_MTM_) vertically along the pylon. The reaction force applied to the bottom of the prosthetic foot is the normal force (F_norm_) relative to the base. For the heel, midfoot, and forefoot axial stiffness testing, the rotatable base was locked at −15°, 0°, and 20° relative to horizontal, respectively. F_norm_ equals F_MTM_/cos(θ). The displacement of the prosthetic foot normal to the base (d_norm_) equals d_MTM_*cos(θ). Therefore, axial prosthetic stiffness (k_pros_) equals the quotient of F_MTM_ and d_MTM_*cos(θ)^2^. We estimated torsional stiffness values from the quotient of the product of F_norm_ and the perpendicular moment arm (r_perp_ = r*cos(θ)) and the angular displacement of the foot 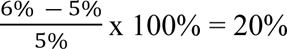).

The displacement of the prosthetic foot normal to the base (d_norm_) equals the product of the vertical displacement of the materials testing machine (d_MTM_) and the cosine of the angle of the base relative to the prosthetic foot (θ) (Fig. 1, 2):

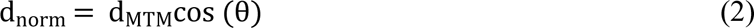

Therefore, the axial stiffness value of the prosthetic foot (k_pros_) equals F_MTM_ divided by the product of d_MTM_ and cos (θ)^2^:

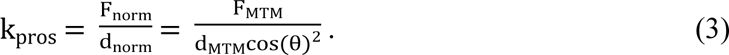

### Torsional Stiffness

We determined torsional stiffness values by dorsiflexing and plantarflexing the prosthetic feet. Plantarflexion and dorsiflexion torsional stiffness values of each prosthetic foot were measured as the quotient of the torque and angular displacement of the prosthesis calculated from the force and displacement measured during the heel and forefoot axial stiffness tests when the rotatable base was locked at −15° and 20°, respectively (Fig. 1). For plantarflexion torsional stiffness, we estimated the moment arm as the horizontal distance between the point of contact of the heel during the heel axial stiffness test and the pylon (r) and multiplied it by cosine of the base angle (−15°) to calculate the perpendicular moment arm (r_perp_). The point of contact was the location where the heel of the prosthesis contacted the base when the prosthetic foot was preloaded with 4-6 N. For dorsiflexion torsional stiffness, we estimated the moment arm as the horizontal distance between the point of contact of the forefoot during the forefoot axial stiffness test and the pylon (r; Fig. 2) and multiplied it by cosine of the base angle (20°) to calculate the perpendicular moment arm (r_perp_; Fig. 2). The point of contact was the location where the forefoot of the prosthesis contacted the base when the prosthetic foot was preloaded with 4-6 N and corresponded with the start of the forefoot axial stiffness test. We calculated torque throughout compression as the product of the normal force (F_norm_) and r_perp_ (Fig. 2). We assumed the point of contact and thus r_perp_ was constant throughout loading and unloading. We calculated the angle of the prosthetic foot (α) as the inverse tangent of the vertical displacement of the MTM (d_MTM_) divided by the horizontal distance of the point of contact and the pylon (r; Fig. 2). Thus, torsional stiffness (*k_pros,torsion_*) equals the quotient of the change in torque (τ) and angle (α in rad) of the prosthetic foot:

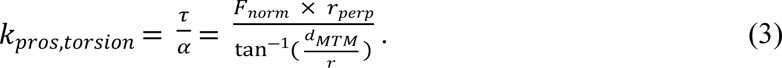

### Hysteresis

We calculated hysteresis for each loading and unloading cycle as the percentage of energy lost during unloading (difference between the energy returned during unloading and the energy stored during loading) compared to the energy stored during loading. Hysteresis was calculated as the quotient of the difference in the area under the loading and unloading curves and the area under the loading curve:

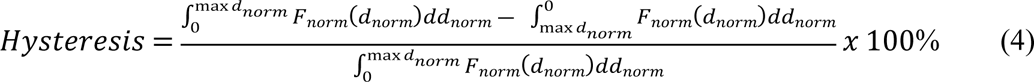

where F_norm_ is the normal force, d_norm_ is the displacement of the prosthetic foot, and dd_norm_ is the differential of the displacement of the prosthetic foot.

### Data Analysis

We used a custom MATLAB script (Mathworks Inc., Natick, MA, USA) to fit linear and quadratic curves to the force-displacement and torque-angle data, calculated average axial and torsional stiffness values, and calculated hysteresis. We used a 20 N F_norm_ threshold to define the start and end of each loading and unloading cycle and set the maximum F_norm_ or torque value of each cycle as the end of the loading phase of the cycle. Then, we fit linear and quadratic least-squares curves to the normal force-displacement and torque-angle data from the loading phases of the last three cycles for each prosthetic foot at the heel, midfoot, and forefoot. We calculated average axial and torsional stiffness values from the discrete value of the slope of the force-displacement and torque-angle curve from the start to the end of the loading phase and averaged that value for the last three test cycles for each foot and test condition. Finally, we averaged the hysteresis from the last three cycles from the normal force-displacement and torque-angle data for each prosthetic foot and test condition.

### Statistical Analysis

We calculated adjusted R^2^ values (27) for the linear and quadratic curves for each prosthetic foot at the heel, midfoot, and forefoot. The axial and torsional stiffness of the prosthetic foot was determined to be better characterized by a linear or quadratic force-displacement or torque-angle curve if the adjusted R^2^ was greater. Then, we constructed eight linear regression models (28) to determine the effect of prosthetic foot stiffness category, prosthetic foot size, and shoe or no shoe on the average axial stiffness values and hysteresis at the heel, midfoot, and forefoot, and torsional stiffness values in the plantarflexion and dorsiflexion directions. We set average axial stiffness values, torsional stiffness values, or hysteresis as the dependent variable and stiffness category (numerical; 1–8), size (numerical; 24– 27 cm), and shoe versus no shoe (categorical; shoe = 1, no shoe = 0) as independent variables. A unit change in hysteresis (%) is a percentage point (p.p.) where one p.p. refers to a 1% unit, such that an increase from 5% to 6% is a 1 p.p. increase as opposed to a 20% increase (i.e. *not* 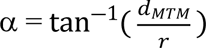). We used a significance level of p < 0.05. All statistical analyses were performed in RStudio (Boston, MA, USA).

## 3. Results

For every prosthetic foot, the adjusted R^2^ was higher when force versus displacement was represented as a quadratic compared to a linear curve (average adjusted R^2^ across all tests– quadratic: 0.99, linear: 0.95). Therefore, the prosthetic foot force-displacement curves were better described by a quadratic compared to a linear fit. The prosthetic foot force-displacement curves were well described by a progressive, quadratic force-displacement curve, meaning that axial stiffness increased with greater force applied (Fig. 3, Fig. 4).

**Fig. 3.**
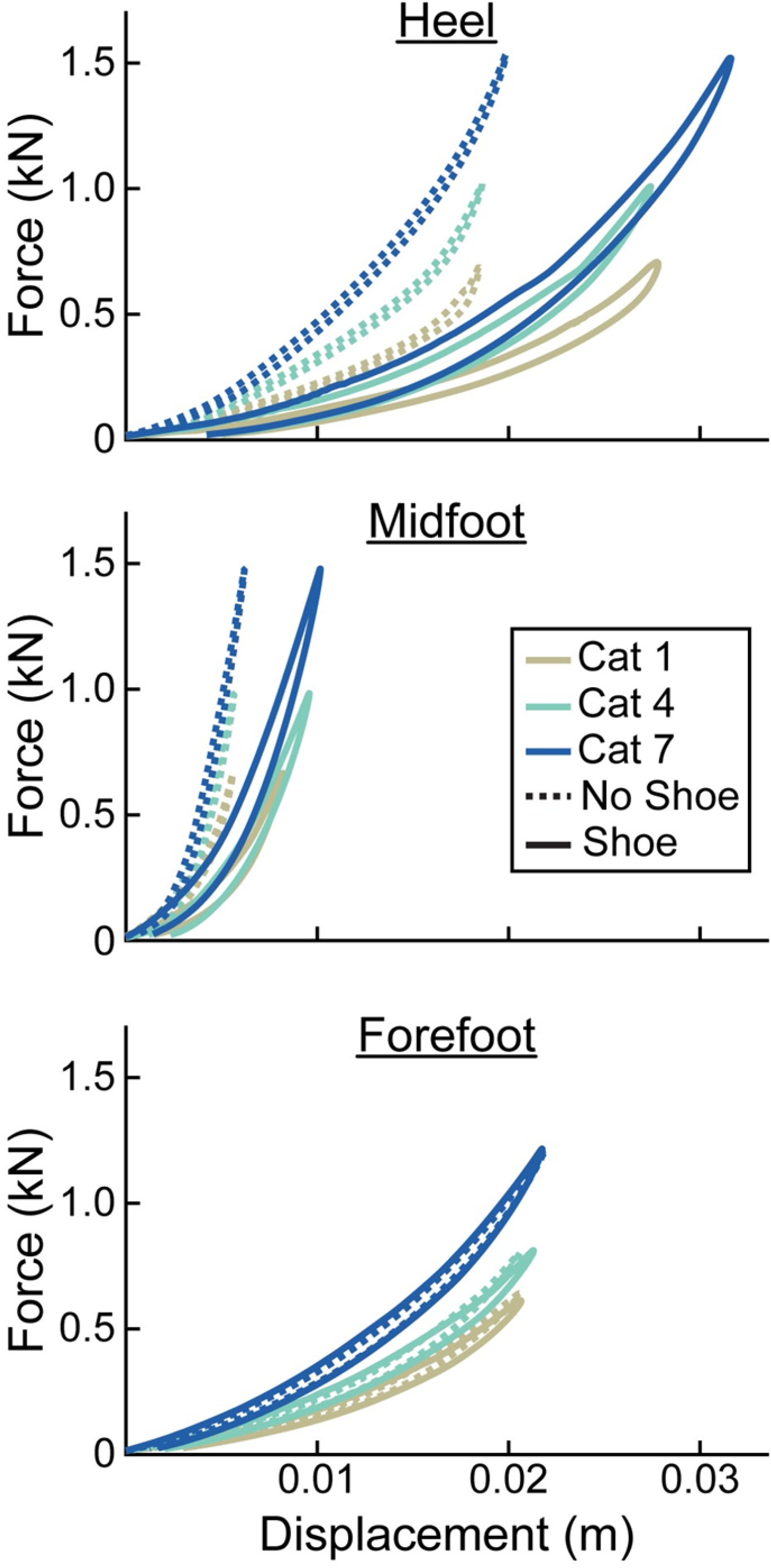
Representative force (kN) versus displacement (m) curves of the heel, midfoot, and forefoot of size 26 cm LP Vari-flex prosthetic feet. The colors represent different stiffness categories (categories 1, 4, 7). The dashed lines are for the tests without a shoe and the solid lines are for the tests with the shoe. Curves go in a clockwise direction from the start to the end of a cycle.

**Fig. 4.**
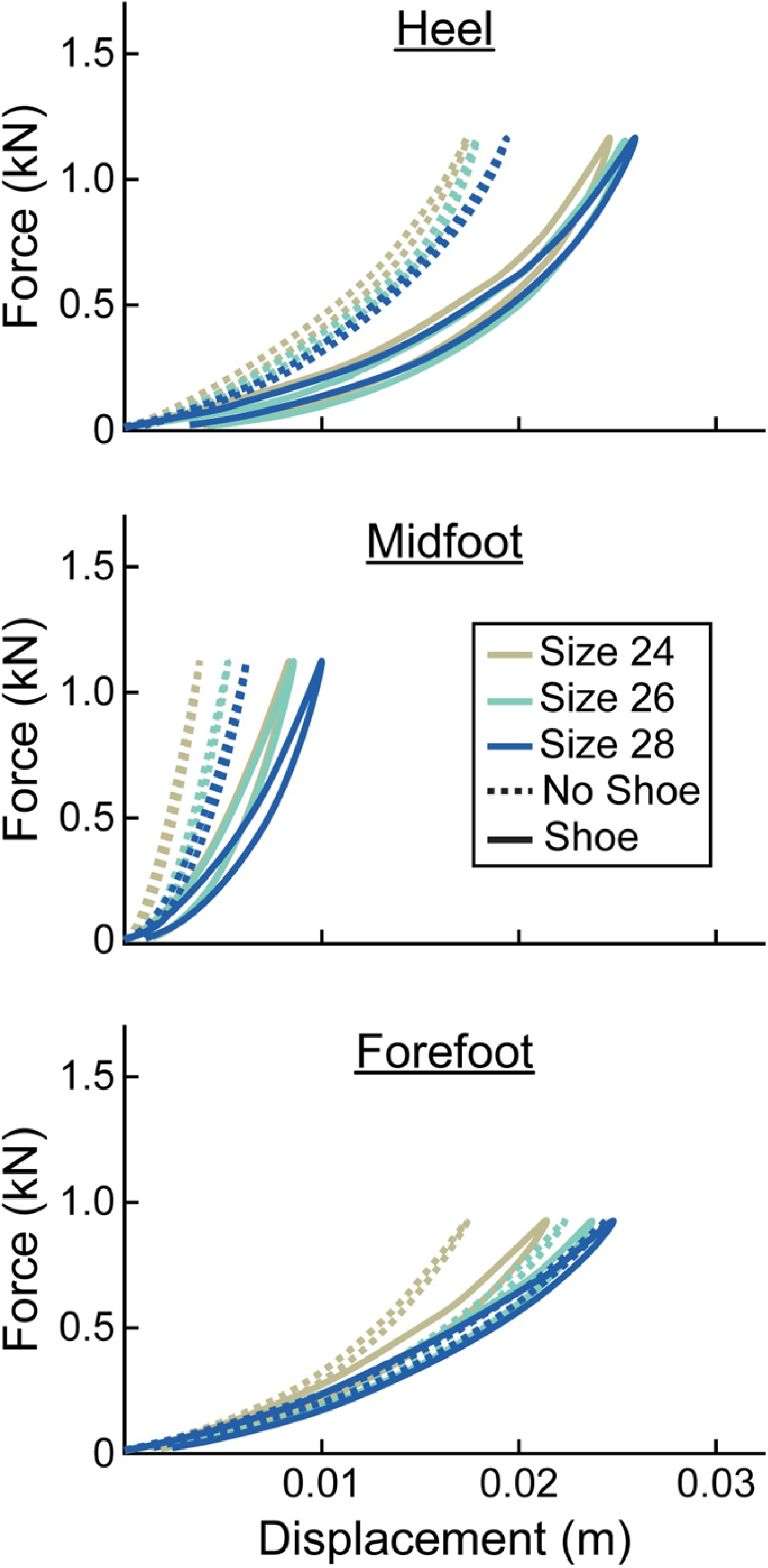
Representative force (kN) versus displacement (m) curves of the heel, midfoot, and forefoot of stiffness category 5 LP Vari-flex prosthetic feet. The colors represent different sizes (24 cm, 26 cm, 28 cm). The dashed lines are for the tests without a shoe and the solid lines are for the tests with the shoe. Curves go in a clockwise direction from the start to the end of a cycle.

At the heel, average prosthetic foot axial stiffness values increased by 5.7 kN/m for every 1 stiffness category increase (p < 0.001), decreased by 1.9 kN/m for every 1 cm increase in size (p < 0.001), and decreased by 14.9 kN/m with the shoe compared to without the shoe (p < 0.001; Fig. 5; Table 2). At the midfoot, average prosthetic foot axial stiffness values increased by 16.0 kN/m for every 1 stiffness category increase (p < 0.001), decreased by 18.9 kN/m for every 1 cm increase in size (p < 0.001), and decreased by 76.2 kN/m with the shoe compared to without the shoe (p < 0.001; Fig. 5; Table 2). At the forefoot, average prosthetic foot axial stiffness values increased by 4.0 kN/m for every 1 stiffness category increase (p < 0.001) and decreased by 1.5 kN/m for every 1 cm increase in size (p < 0.001; Fig. 5; Table 2). However, we did not detect a statistically significant effect of shoe on the average forefoot prosthetic foot axial stiffness value (p = 0.27; Fig. 5; Table 2).

**Fig. 5.**
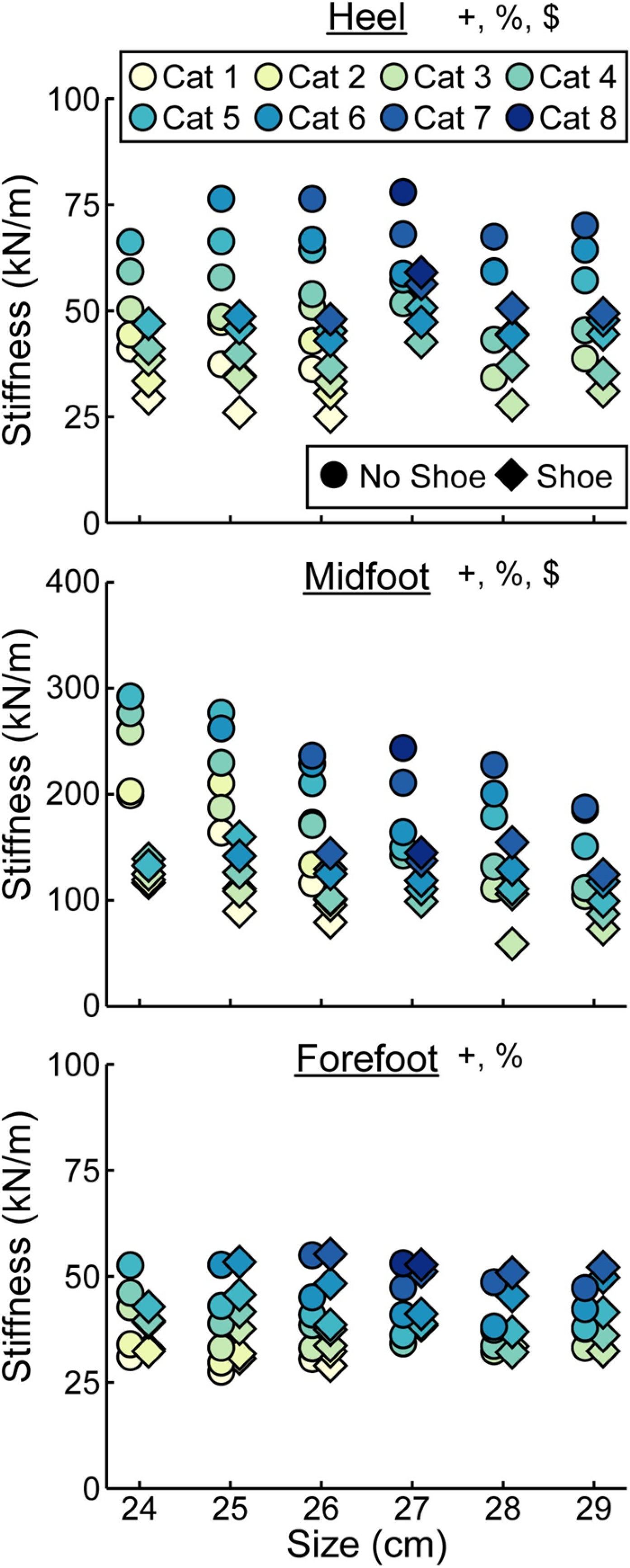
**Average axial stiffness values (kN/m) versus LP Vari-flex prosthetic foot size in cm.** The colors represent different stiffness categories (categories 1–8), the circles represent average axial stiffness values without a shoe, and the diamonds represent average axial stiffness values with a shoe. Symbols are offset for no shoe and shoe for clarity. The y-axis differs for the midfoot compared to heel and forefoot axial stiffness values. + indicates a significant effect of stiffness category, % indicates a significant effect of size, and $ indicates a significant effect of shoe.

**Fig. 6.**
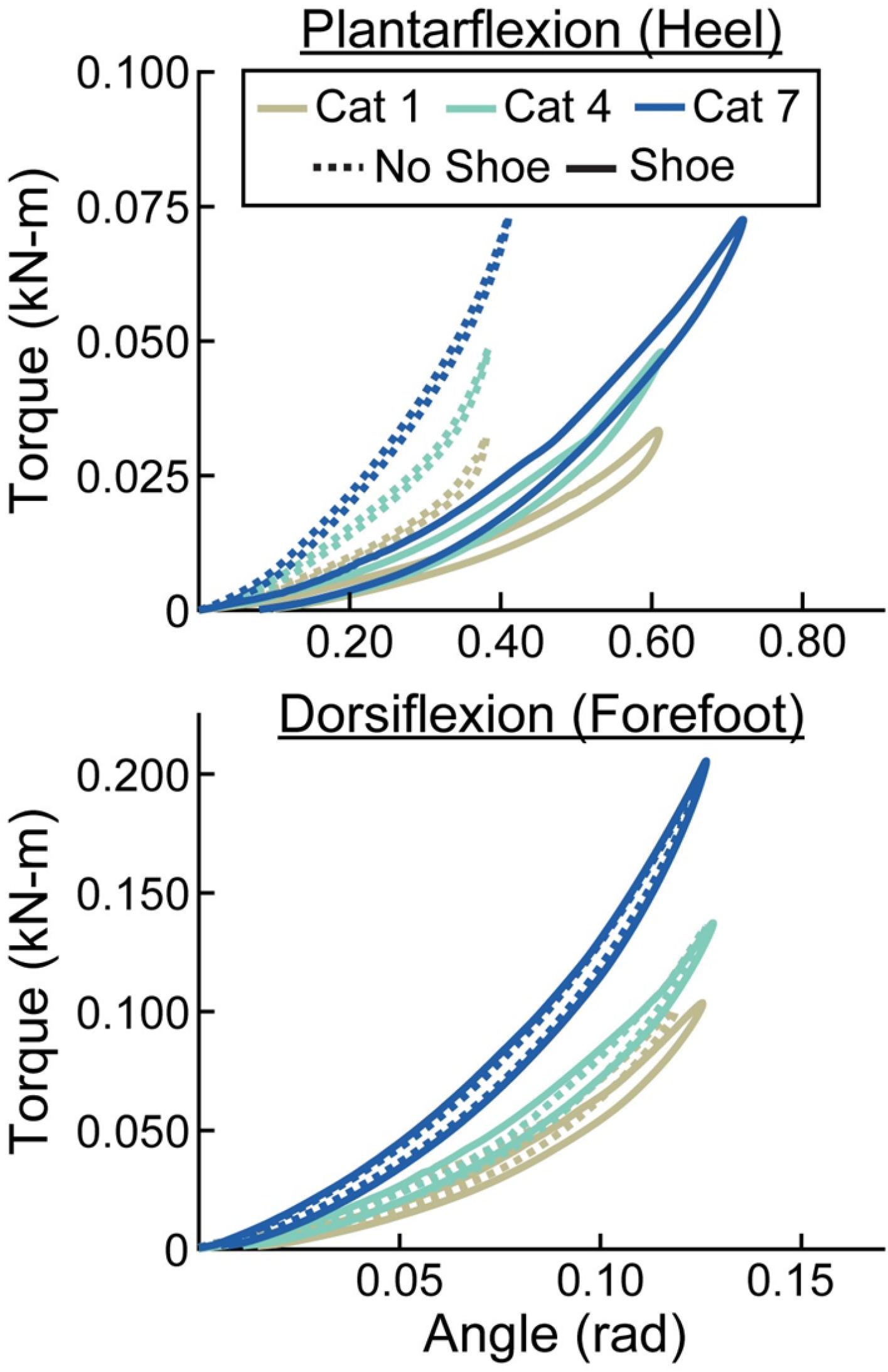
Representative torque (kN-m) versus angle (rad) curves for plantarflexion (heel) and dorsiflexion (forefoot) of size 26 cm LP Vari-flex prosthetic feet. The colors represent different stiffness categories (categories 1, 4, 7). The dashed lines are for the tests without a shoe and the solid lines are for the tests with the shoe. The x- and y-axes differ for the plantarflexion and dorsiflexion torque and angle values.

**Fig. 7.**
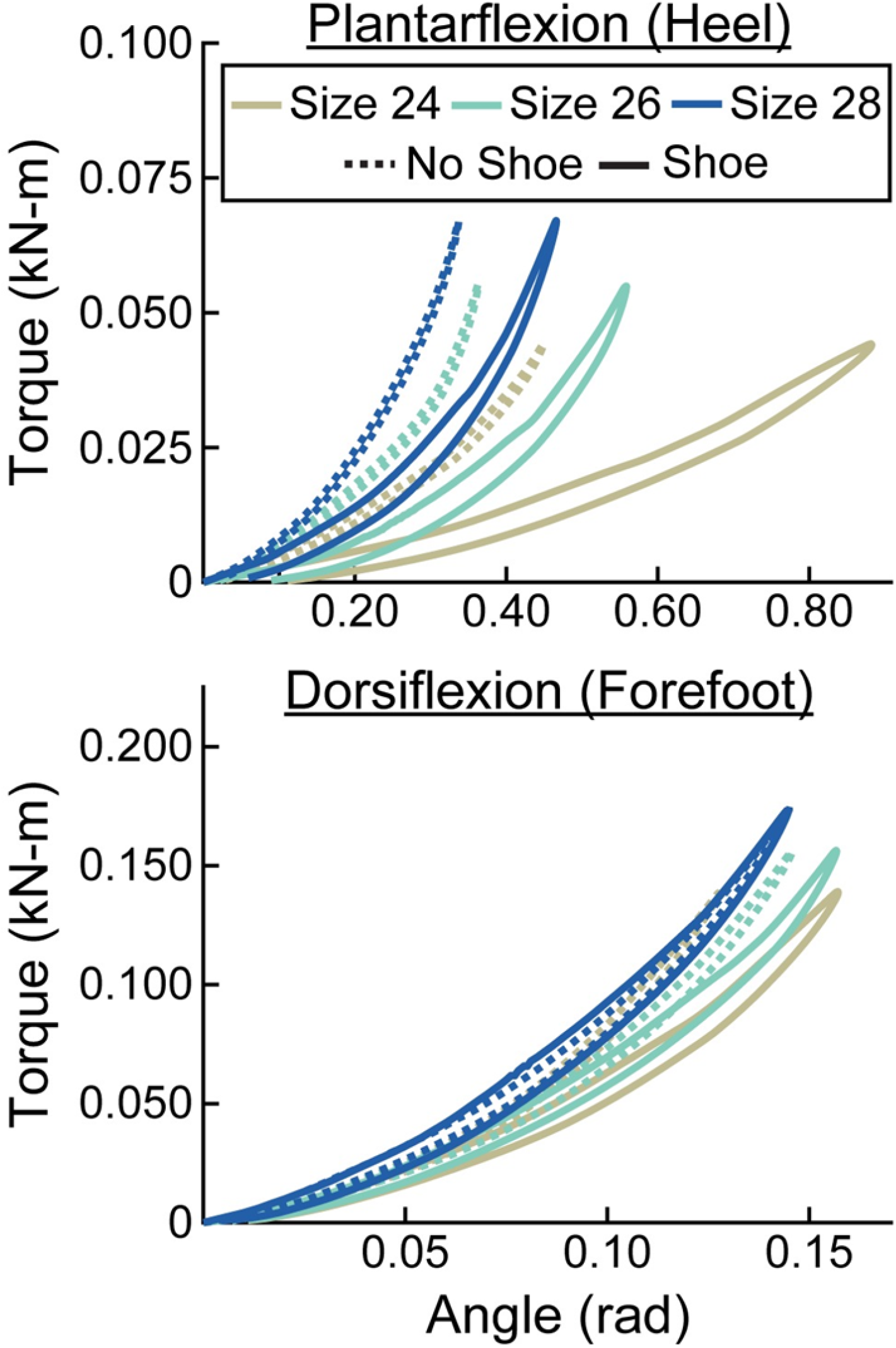
Representative torque (kN-m) versus angle (rad) curves for plantarflexion (heel) and dorsiflexion (forefoot) of category 5 stiffness LP Vari-flex prosthetic feet. The colors represent different sizes (24 cm, 26 cm, 28 cm). The dashed lines are for the tests without a shoe and the solid lines are for the tests with the shoe. The x- and y-axes differ for the plantarflexion and dorsiflexion torque and angle values.

**Table 2.**
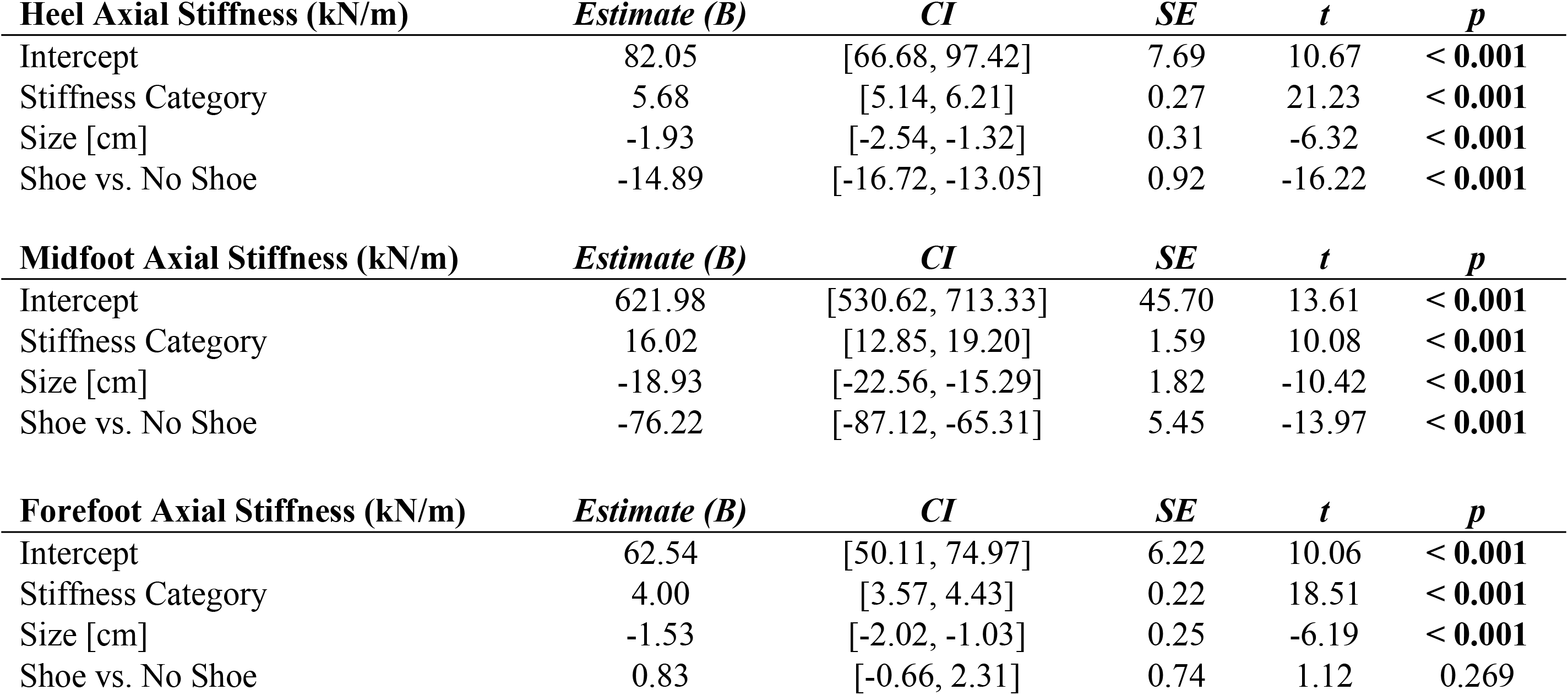
Linear regression parameters for fixed effects of LP Vari-flex prosthetic foot stiffness category, size, and shoe or no shoe on the axial stiffness values (kN/m) at the heel, midfoot, and forefoot. Coefficient estimates, 95% confidence intervals for coefficient estimates (CI), coefficient standard errors (SE), t values (t), and p values (p) are listed for each stiffness category (1–8) and size (24–29 cm). The shoe vs. no shoe coefficient is in reference to the no shoe condition.

| **Heel Axial Stiffness (kN/m)** | *Estimate (B)* | *CI* | *SE* | *t* | *p* |
| --- | --- | --- | --- | --- | --- |
| Intercept | 82.05 | [66.68, 97.42] | 7.69 | 10.67 | < **0.001** |
| Stiffness Category | 5.68 | [5.14, 6.21] | 0.27 | 21.23 | < **0.001** |
| Size [cm] | -1.93 | [-2.54, -1.32] | 0.31 | -6.32 | < **0.001** |
| Shoe vs. No Shoe | -14.89 | [-16.72, -13.05] | 0.92 | -16.22 | < **0.001** |
| **Midfoot Axial Stiffness (kN/m)** | *Estimate (B)* | *CI* | *SE* | *t* | *p* |
| Intercept | 621.98 | [530.62, 713.33] | 45.70 | 13.61 | < **0.001** |
| Stiffness Category | 16.02 | [12.85, 19.20] | 1.59 | 10.08 | < **0.001** |
| Size [cm] | -18.93 | [-22.56, -15.29] | 1.82 | -10.42 | < **0.001** |
| Shoe vs. No Shoe | -76.22 | [-87.12, -65.31] | 5.45 | -13.97 | < **0.001** |
| **Forefoot Axial Stiffness (kN/m)** | *Estimate (B)* | *CI* | *SE* | *t* | *p* |
| Intercept | 62.54 | [50.11, 74.97] | 6.22 | 10.06 | < **0.001** |
| Stiffness Category | 4.00 | [3.57, 4.43] | 0.22 | 18.51 | < **0.001** |
| Size [cm] | -1.53 | [-2.02, -1.03] | 0.25 | -6.19 | < **0.001** |
| Shoe vs. No Shoe | 0.83 | [-0.66, 2.31] | 0.74 | 1.12 | 0.269 |

When force was applied at the heel, average prosthetic foot plantarflexion torsional stiffness values increased by 0.01 kN-m/rad for every 1 stiffness category increase (p < 0.001), increased by 0.02 kN-m/rad for every 1 cm increase in size (p < 0.001), and decreased by 0.05 kN-m/rad with the shoe compared to without the shoe (p < 0.001; Fig. 8; Table 3). When force was applied at the forefoot, average prosthetic foot dorsiflexion torsional stiffness values increased by 0.13 kN-m/rad for every 1 stiffness category increase (p < 0.001) and increased by 0.09 kN-m/rad for every 1 cm increase in size (p < 0.001; Fig. 8; Table 3). However, we did not detect a statistically significant effect of shoe on the average prosthetic foot dorsiflexion torsional stiffness value (p = 0.19; Fig. 8; Table 3).

**Fig. 8.**
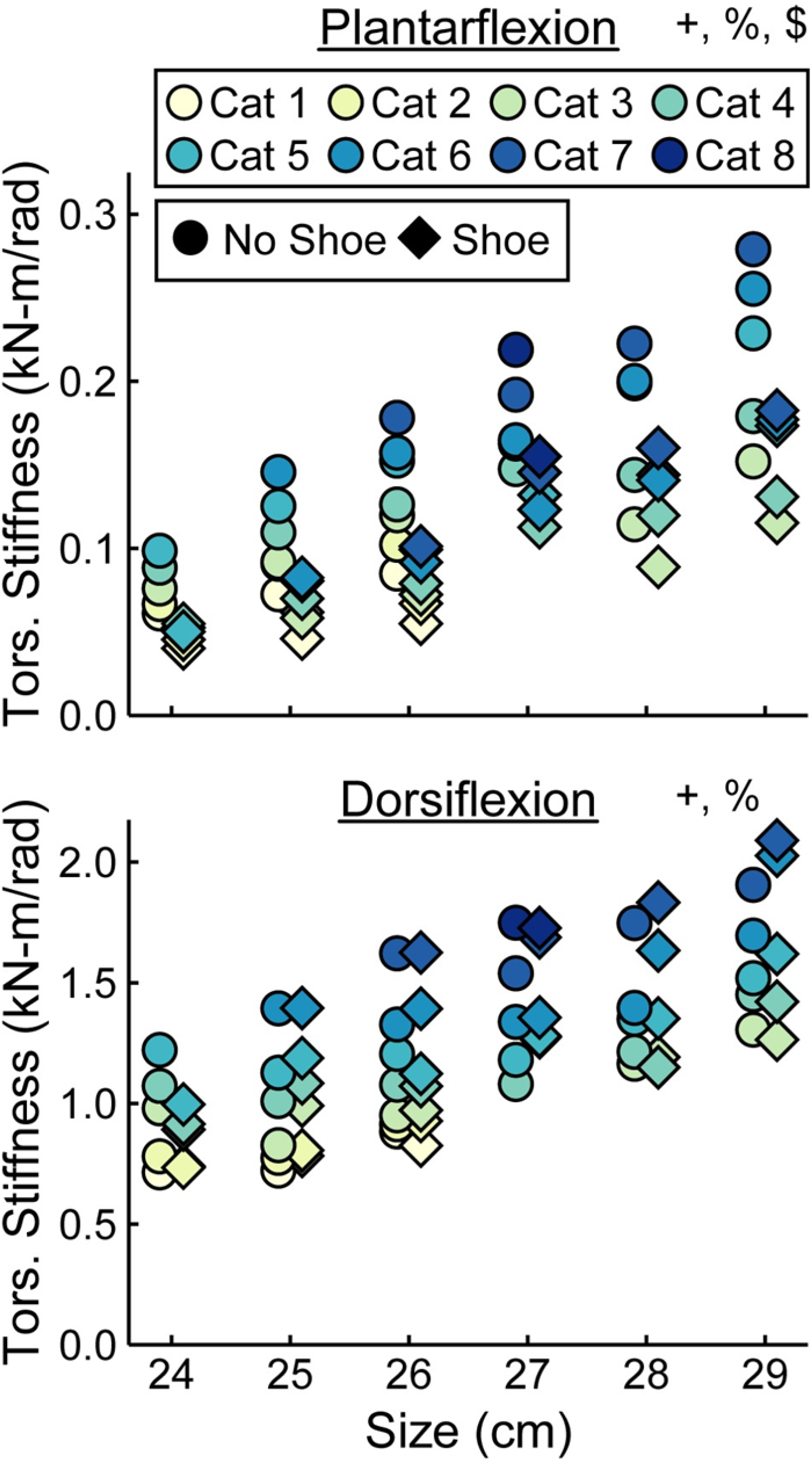
Average torsional (tors.) stiffness values (kN-m/rad) versus LP Vari-flex prosthetic foot size in cm for plantarflexion (heel) and dorsiflexion (forefoot). The colors represent different stiffness categories (categories 1–8), the circles represent average torsional stiffness without a shoe, and the diamonds represent average torsional stiffness with a shoe. Symbols are offset for no shoe and shoe for clarity. The y-axis differs for the plantarflexion and dorsiflexion torsional stiffness values. + indicates a significant effect of stiffness category, % indicates a significant effect of size, and $ indicates a significant effect of shoe.

**Table 3.** Linear regression parameters for fixed effects of LP Vari-flex prosthetic foot stiffness category, size, and shoe or no shoe on the torsional stiffness values (kN-m/rad) in plantarflexion (heel) and dorsiflexion (forefoot). Coefficient estimates, 95% confidence intervals for coefficient estimates (CI), coefficient standard errors (SE), t values (t), and p values (p) are listed for each stiffness category (1–8) and size (24–29 cm). The shoe vs. no shoe coefficient is in reference to the no shoe condition.

| **Plantarflexion (Heel) Torsional Stiffness (kN-m/rad)** | *Estimate (B)* | *CI* | *SE* | *t* | *p* |
| --- | --- | --- | --- | --- | --- |
| Intercept | -0.40 | [-0.47, -0.33] | 0.03 | -11.67 | < **0.001** |
| Stiffness Category | 0.01 | [0.01, 0.02] | 0.00 | 11.46 | < **0.001** |
| Size [cm] | 0.02 | [0.02, 0.02] | 0.00 | 13.54 | < **0.001** |
| Shoe vs. No Shoe | -0.05 | [-0.05, -0.04] | 0.00 | -11.34 | < **0.001** |
| **Dorsiflexion (Forefoot) Torsional Stiffness (kN-m/rad)** | *Estimate (B)* | *CI* | *SE* | *t* | *p* |
| Intercept | -1.61 | [-2.05, -1.18] | 0.22 | -7.41 | < **0.001** |
| Stiffness Category | 0.13 | [0.11, 0.14] | 0.01 | 16.72 | < **0.001** |
| Size [cm] | 0.09 | [0.07, 0.10] | 0.01 | 9.97 | < **0.001** |
| Shoe vs. No Shoe | 0.03 | [-0.02, 0.09] | 0.03 | 1.32 | 0.192 |

Hysteresis at the heel decreased by 0.3 percentage points (p.p.) for every 1 stiffness category increase (p < 0.001), decreased by 1.0 p.p. for every 1 cm increase in size (p = 0.01), and increased by 13.8 p.p. with the shoe compared to without the shoe (p < 0.001; Fig. 9, Table 4). Hysteresis at the midfoot decreased by 0.3 p.p. for every 1 stiffness category increase (p = 0.04), decreased by 0.5 p.p. for every 1 cm increase in size (p = 0.01), and increased by 11.0 p.p. with the shoe compared to without the shoe (p < 0.001; Fig. 9, Table 4). Hysteresis at the forefoot decreased by 0.3 p.p. for every 1 stiffness category increase (p = 0.01) and increased by 7.0 p.p. with the shoe compared to without the shoe (p < 0.001; Fig. 9, Table 4). However, we did not detect a statistically significant effect of prosthetic foot size on hysteresis at the forefoot (p = 0.48; Fig. 9, Table 4).

**Fig. 9.**
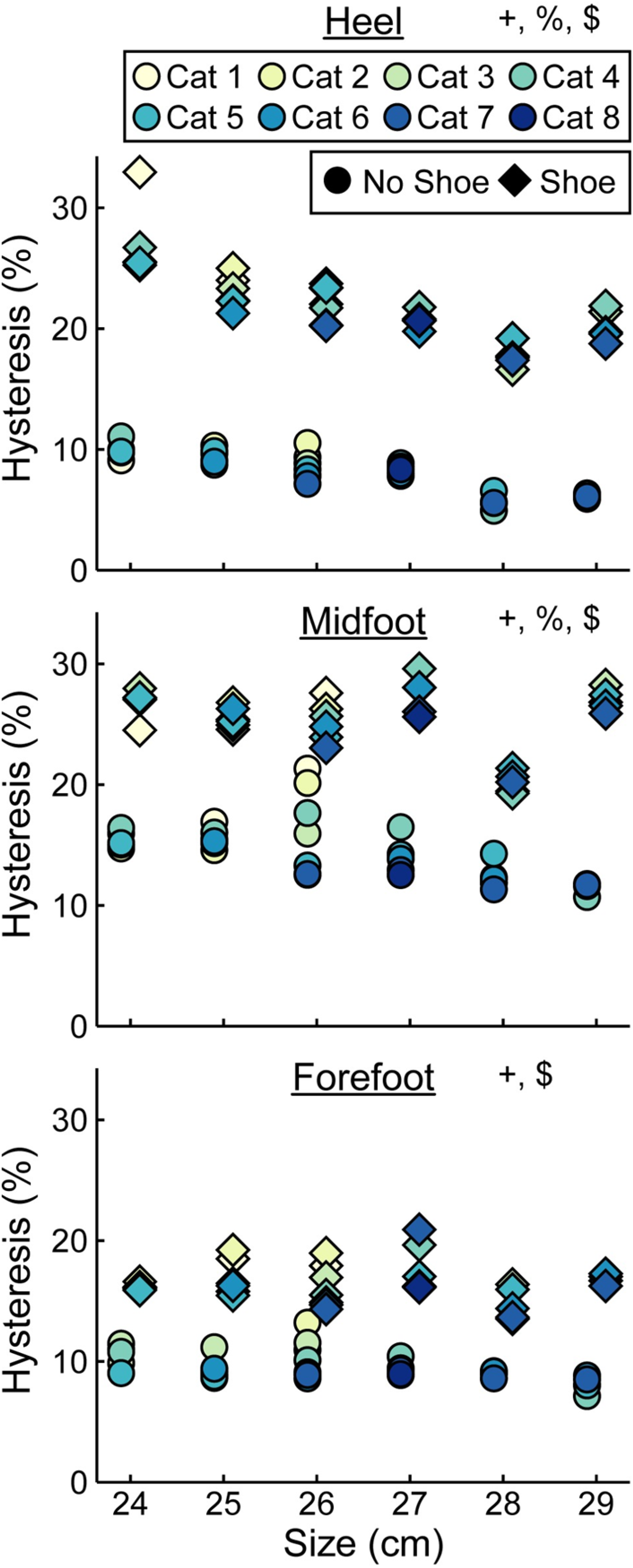
**Average hysteresis (%) versus LP Vari-flex prosthetic foot size in cm.** The colors represent different prosthetic foot stiffness categories (categories 1–8), the circles represent average hysteresis without a shoe, and the diamonds represent average hysteresis with a shoe. Symbols are offset for no shoe and shoe for clarity. + indicates a significant effect of stiffness category, % indicates a significant effect of size, and $ indicates a significant effect of shoe.

**Table 4.** Linear regression parameters for fixed effects of LP Vari-flex prosthetic foot stiffness category, size, and shoe or no shoe on the hysteresis (%) at the heel, midfoot, and forefoot. Coefficient estimates, 95% confidence intervals for coefficient estimates (CI), coefficient standard errors (SE), t values (t), and p values (p) are listed for each stiffness category (1–8) and size (24–29 cm). The shoe vs. no shoe coefficient is in reference to the no shoe condition.

| **Heel Hysteresis (%)** | ***Estimate (B)*** | ***CI*** | ***SE*** | ***t*** | ***p*** |
| --- | --- | --- | --- | --- | --- |
| Intercept | 36.56 | [29.95, 43.17] | 3.31 | 11.06 | < **0.001** |
| Stiffness Category | -0.31 | [-0.54, -0.08] | 0.12 | -2.70 | **0.009** |
| Size [cm] | -1.03 | [-1.29, -0.76] | 0.13 | -7.81 | < **0.001** |
| Shoe vs. No Shoe | 13.81 | [13.02, 14.60] | 0.39 | 35.00 | < **0.001** |
| **Midfoot Hysteresis (%)** | ***Estimate (B)*** | ***CI*** | ***SE*** | ***t*** | ***p*** |
| Intercept | 28.40 | [18.96, 37.84] | 4.72 | 6.01 | < **0.001** |
| Stiffness Category | -0.34 | [-0.66, -0.01] | 0.16 | -2.05 | **0.045** |
| Size [cm] | -0.48 | [-0.85, -0.10] | 0.19 | -2.54 | **0.014** |
| Shoe vs. No Shoe | 10.99 | [9.86, 12.12] | 0.56 | 19.49 | < **0.001** |
| **Forefoot Hysteresis (%)** | ***Estimate (B)*** | ***CI*** | ***SE*** | ***t*** | ***p*** |
| Intercept | 12.78 | [7.22, 18.34] | 2.78 | 4.60 | < **0.001** |
| Stiffness Category | -0.28 | [-0.47, -0.08] | 0.10 | -2.85 | **0.006** |
| Size [cm] | -0.08 | [-0.30, 0.14] | 0.11 | -0.71 | 0.479 |
| Shoe vs. No Shoe | 6.99 | [6.32, 7.65] | 0.33 | 21.05 | < **0.001** |

## 4. Discussion

In support of our first hypothesis, the force-displacement curves of the LP Vari-flex prosthetic foot at the heel, midfoot, and forefoot (Supplementary Material: Force-Displacement Equations) and the plantarflexion and dorsiflexion torque-angle curves (Supplementary Material: Torque-Angle Equations) exhibited a curvilinear profile and were well-described by a quadratic curve (average adjusted R^2^ for all tests: 0.99). The LP Vari-flex prosthetic foot exhibits a curvilinear force-displacement profile like its higher profile version, the Vari-flex (9), likely because of the similar design and because both prostheses are made of carbon fiber. The LP Vari-flex prosthetic foot force-displacement and torque-angle curves have a steeper slope with greater applied forces and torques and thus stiffens with displacement, which suggests that the stiffness of the prosthetic foot may vary between different tasks.

In support of our second hypothesis that a greater prosthetic foot stiffness category would result in increased axial stiffness values, we found that a greater numerical stiffness category of the LP Vari-flex prosthetic foot resulted in increased average axial stiffness values at the heel, midfoot, and forefoot by 5.8, 16.0, and 4.0 kN/m per one category increase, respectively. The change in heel stiffness values between categories of the LP Vari-flex prosthetic foot is similar to the higher profile version of the prosthetic foot (Vari-flex), which has a heel axial stiffness value that increases by about 6–7 kN/m per one category increase (9,17). In general, the heel axial stiffness value of the LP Vari-flex prosthetic foot for a given stiffness category is stiffer than the higher profile Vari-flex prosthetic foot. For example, when comparing prosthetic foot stiffness categories and sizes common to this study and other studies that characterized Vari-flex prosthetic feet, the average axial stiffness value of the heel for the size 27 LP Vari-flex and Vari-flex prosthetic feet without shoes for categories 5–7 range from 57.3–68.1 kN/m and 36.4–59.2 kN/m, respectively (9,17). Moreover, the forefoot axial stiffness value of the LP Vari-flex prosthetic foot for a given category is stiffer than the forefoot stiffness measured for the Vari-flex prosthetic foot. For example, the average axial stiffness value of the forefoot for the size 27 LP Vari-flex and Vari-flex prosthetic feet without shoes for categories 5–7 range from 36.1–47.4 kN/m and 28.6–40.0 kN/m, respectively (9). Ultimately, the differences in heel and forefoot axial stiffness values between the LP Vari-flex and Vari-flex prosthetic feet suggest that prosthetists should prescribe a lower stiffness category for the LP Vari-flex prosthesis than they would for the Vari-flex prosthesis. Furthermore, the changes in axial stiffness values of the LP Vari-flex prosthetic foot without a shoe between prosthetic stiffness categories for the heel, midfoot, and forefoot were variable (Fig. 5). Previous studies that characterized commercially available passive-elastic prosthetic feet found similar results (9,17). The variable changes in axial stiffness values between stiffness categories highlight the need for objective measurements of prosthetic foot stiffness within and between manufacturers.

In further support of our second hypothesis that a greater prosthetic foot stiffness category would result in an increased torsional stiffness value, we found that a greater stiffness category of the LP Vari-flex prosthetic foot resulted in an increased average torsional stiffness value for plantarflexion (heel) and dorsiflexion (forefoot) by 0.01 and 0.13 kN-m/rad, respectively. Major et al. estimated the torsional stiffness values of three commercially-available prosthetic feet, the SACH foot, Seattle foot, and Flex-foot, and found that the plantarflexion (heel) and dorsiflexion (forefoot) stiffness values ranged from 0.09–0.20 kN-m/rad and 0.39–1.40 kN-m/rad, respectively (6). Similarly, we found that plantarflexion (heel) stiffness values of the LP Vari-flex foot without a shoe ranged from 0.06–0.28 kN-m/rad and dorsiflexion (forefoot) stiffness values of the LP Vari-flex foot without a shoe ranged from 0.71–1.91 kN-m/rad across the tested stiffness categories and sizes. Major et al. found that different torsional stiffness values of prosthetic feet can affect the joint angles, peak ground reaction forces, and metabolic cost of people with unilateral transtibial amputation during walking (6), so characterizing the torsional stiffness values of prosthetic feet can be a useful tool for predicting how different prosthetic feet will affect walking biomechanics. Moreover, since a biological ankle can behave mechanically like a torsional spring and damper system during walking at 1.2 m/s (19), torsional stiffness values of prosthetic feet provide information that can be compared to the biological ankle-foot system (29) to derive function and potentially inform biomimetic prosthetic prescription and design.

In contrast to our second hypothesis that a greater prosthetic foot stiffness category would not affect hysteresis with or without a shoe, we found that a greater LP Vari-flex prosthetic foot stiffness category resulted in a 0.3 percentage point decrease in hysteresis for the heel, midfoot, and forefoot. Even though we found a significant effect of prosthetic foot stiffness category on hysteresis, the effects on hysteresis at the heel, midfoot, and forefoot were small. The hysteresis at the heel, midfoot, and forefoot of the LP Vari-flex prosthetic feet without a shoe averaged across sizes ranged from 6.9–10.3%, 12.1–18.1%, and 8.8–10.7%, respectively. Therefore, it is unclear if the effect of LP Vari-flex prosthetic foot stiffness category on hysteresis is clinically meaningful. Nonetheless, prosthetists may want to consider that LP Vari-flex prosthetic feet with stiffer categories have less hysteresis than less stiff categories when prescribing prosthetic feet.

We partially reject our third hypothesis that an increase in prosthetic foot size would have no effect on axial stiffness values or hysteresis but would increase torsional stiffness values. A 1 cm increase in the size of LP Vari-flex prosthetic feet resulted in a decrease of the axial stiffness values at the heel, midfoot, and forefoot by 1.9, 18.9, and 1.5 kN/m, respectively, an increase in plantarflexion (heel) and dorsiflexion (forefoot) torsional stiffness values by 0.02 and 0.09 kN-m/rad, respectively, and a decrease in the hysteresis at the heel and midfoot by 1.0 and 0.5 percentage points, respectively. As hypothesized, an increase in prosthetic foot size resulted in an increase in torsional stiffness values due to an increase in the moment arm of the prosthesis. Despite the fact that manufacturers recommend the same LP Vari-flex prosthetic foot stiffness category for a given body mass and activity level regardless of prosthetic foot size (8), prosthetic foot size does affect axial stiffness values, torsional stiffness values, and hysteresis. Since axial stiffness values, torsional stiffness values, and hysteresis can affect kinematics, kinetics, muscle activity, metabolic cost and user preference during walking (1–7), prosthetists should consider that an increase in the size of the LP Vari-flex prosthetic foot can decrease axial stiffness values, decrease hysteresis, and increase torsional stiffness values when prescribing prosthetic feet. Furthermore, prosthetic feet should be designed to have similar mechanical properties for a given prosthetic foot stiffness category regardless of the prosthetic foot size.

In support of our fourth hypothesis that adding a shoe to the LP Vari-flex prosthetic foot would lower the heel and midfoot axial stiffness values and increase hysteresis, we found that adding a shoe decreased axial stiffness values at the heel and midfoot by 14.9 and 76.2 kN/m, respectively, and increased hysteresis at the heel, midfoot, and forefoot by 13.8, 11.0, and 7.0 percentage points, respectively. Our results are similar to those of Major et al. who found that adding an athletic shoe to the prosthetic foot decreased heel and midfoot axial stiffness values by 20.5 kN/m and 151.6 kN/m, respectively, and increased hysteresis at the heel, midfoot, and forefoot by 7.4, 9.3, and 3.4 percentage points, respectively, compared to values for a prosthetic foot without a shoe (23). Moreover, similar to Major et al., we found that adding a shoe did not affect forefoot axial stiffness values (23). Ultimately, adding a shoe to a prosthetic foot affects the heel and midfoot axial stiffness values and heel, midfoot, and forefoot hysteresis, so footwear should be considered when determining how different prosthetic feet affect kinematics, kinetics, muscle activity, metabolic cost, and user preference of people with transtibial or transfemoral amputation.

In contrast to our fourth hypothesis that adding a shoe to the LP Vari-flex prosthetic foot would not affect torsional stiffness values, we found that adding a shoe resulted in a decrease of plantarflexion (heel) torsional stiffness by 0.05 kN-m/rad but did not affect dorsiflexion (forefoot) torsional stiffness. Adding a shoe to the LP Vari-flex prosthetic foot affects heel and midfoot axial stiffness values, heel, midfoot, and forefoot hysteresis, and plantarflexion (heel) torsional stiffness values. Previous studies have found that different types of footwear can have different effects on stiffness and hysteresis (23). This highlights the need to consider footwear when prescribing prosthetic feet and predicting how different prosthetic feet may affect kinematics, kinetics, muscle activity, metabolic cost, and user preference of people with transtibial or transfemoral amputation during walking.

Our study had some potential limitations. We used a uniaxial load cell (Instron 2580-201, Norwood, MA), so we were unable to measure off-axis forces on the load cell during the heel and forefoot tests. We used a low-friction roller system to reduce off-axis forces on the load cell and derived equations (1)-(3) to estimate the actual force applied to the prosthetic foot based on the force measured by the uniaxial load cell (Supplementary Material: Derivation and Verification of Equations 1–3). However, since the low-friction roller system is not perfectly frictionless, we may have overestimated the force on the prosthetic foot (Supplementary Material: Derivation and Verification of Equations 1–3). We conducted a post-hoc analysis of the forefoot test with one prosthetic foot using a multi-axis force transducer (MC3A-500, AMTI, Watertown, MA, USA) and found that our estimate of the force applied to the prosthetic foot from equation (1) overestimated the actual force measured by the multi-axis force transducer by 1% (Supplementary Material: Derivation and Verification of Equations 1–3). Another potential limitation is that we estimated the torque and angle of each prosthetic foot assuming a constant moment arm (r) from the point of contact of the foot to the pylon when the prosthesis was preloaded to 4–6 N for the heel and forefoot tests. However, r likely decreases as the prosthetic foot is plantarflexed during the heel test and dorsiflexed during the forefoot test. Therefore, we may have overestimated the torque on the prosthetic foot.

## 5. Conclusions

We characterized the axial stiffness values, torsional stiffness values, and hysteresis of LP Vari-flex prosthetic feet with a range of stiffness categories and sizes without and with shoes. In general, a greater prosthetic foot stiffness category resulted in an increase in heel, midfoot, and forefoot axial stiffness values, an increase in plantarflexion and dorsiflexion torsional stiffness values, and a decrease in heel, midfoot, and forefoot hysteresis. Moreover, an increase in prosthetic foot size resulted in a decrease in heel, midfoot, and forefoot axial stiffness values, an increase in plantarflexion and dorsiflexion torsional stiffness values, and a decrease in heel and midfoot hysteresis. Finally, adding a shoe to the LP Vari-flex prosthetic foot resulted in a decrease in heel and midfoot axial stiffness values, a decrease in plantarflexion torsional stiffness values, and an increase in heel, midfoot, and forefoot hysteresis.

Overall, the axial and torsional stiffness values, hysteresis, and force-displacement equations of LP Vari-flex prosthetic feet with and without a shoe can be used to objectively compare LP Vari-flex prosthetic feet to other prosthetic feet in order to inform their prescription and design and use by people with a transtibial or transfemoral amputation. For example, prosthetists can compare our objective stiffness values to values reported for other prosthetic feet (9) rather than using manufacturer-defined categories that can be inconsistent or subjective. In addition, our results can be used by researchers conducting studies on the effects of different prosthetic devices to gain a better understanding of the mechanical properties that affect walking kinematics, kinetics, muscle activity, metabolic cost, and user preference.

